## Supplemental table 1-4, supplemental figure 1-7 for "Maternal age modulates progeny social behavior via a small RNA–neuropeptide axis"

### Supplemental Figure, Table and Legends

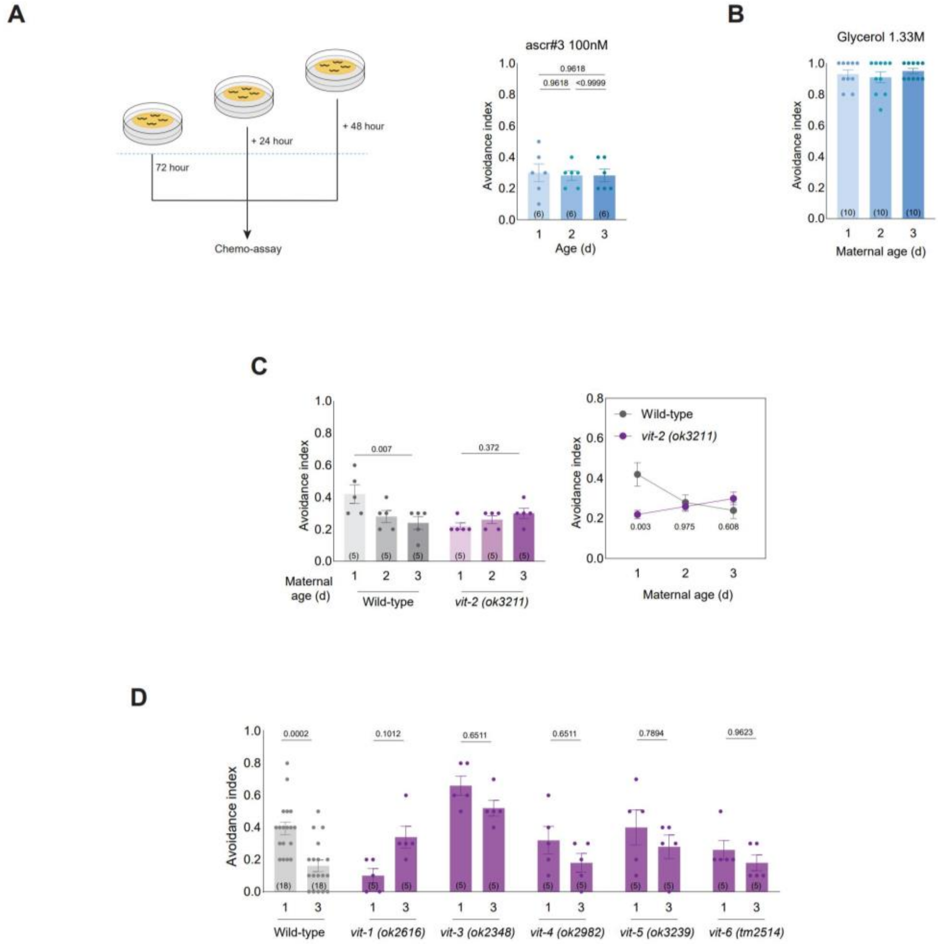


**Supplemental Figure S1. Maternal age shapes progeny avoidance of ascr#3.** (A) Schematic of the behavioral assay with different ages. Synchronized wild-type animals of different ages were tested on the same day. Avoidance index of wild-type animals with different ages with 100 nM ascr#3. *N* = 6. (B) Avoidance index of wild-type animals with different maternal ages with 1.33 M glycerol. *N* = 10. (C) Avoidance index of wild-type animals and *vit-2 (ok3211)* mutants with different maternal ages with 100 nM ascr#3. *N* = 5. (D) Avoidance index of wild-type animals and *vit-1 (ok2616), vit-3 (ok2348), vit-4 (ok2982), vit-5 (ok3239)*, and *vit-6 (tm2514)* mutants with different maternal ages with 100 nM ascr#3. *N* = 5–18. Error bars indicate SEM. One-way ANOVA followed by Tukey's multiple comparisons (A–C); two-way AVOVA followed by Sidak's multiple comparisons (D).

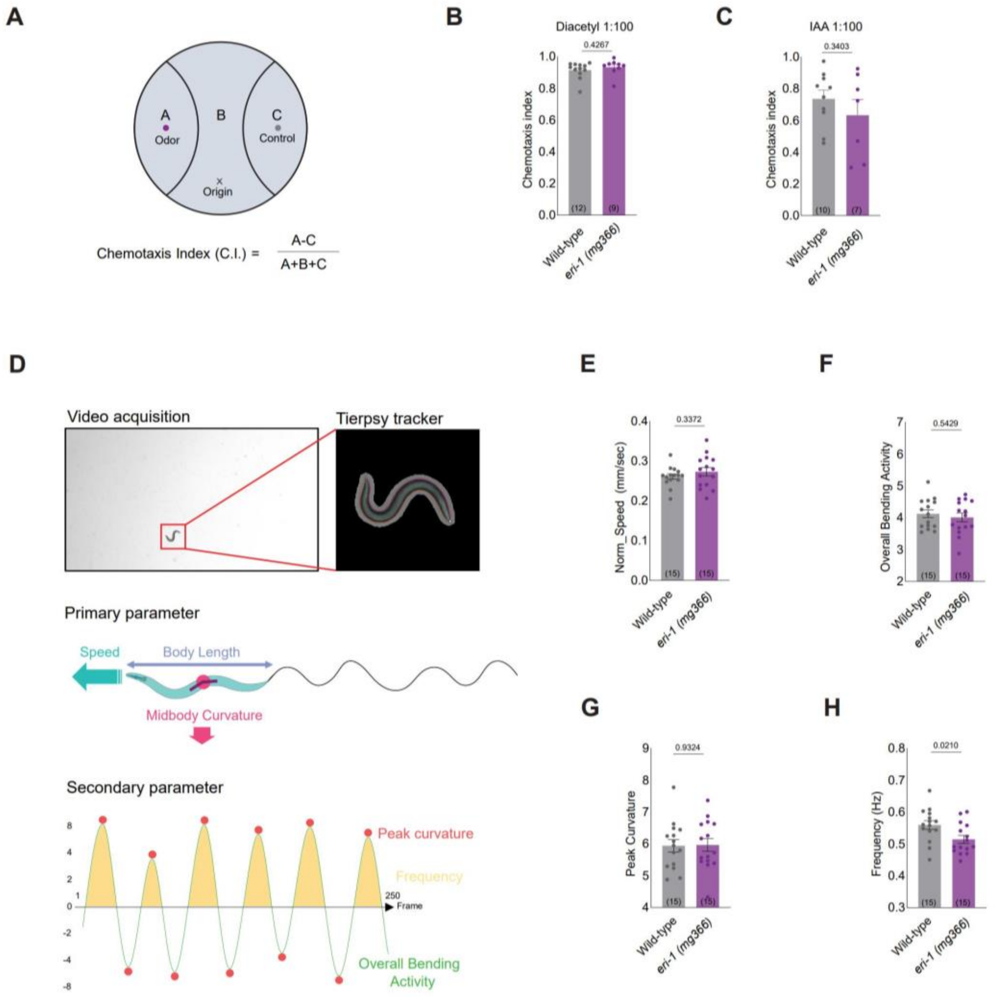


**Supplemental Figure S2.** ***eri-1*** **mutants show normal attraction and locomotion behaviors.** (A) Schematic of the chemotaxis assay (see Star Methods). (B) Chemotaxis index of wild-type animals and *eri-1 (mg366)* mutants in response to 1:100 diacetyl. $N$ = 9–12. (C) Chemotaxis index of wild-type animals and *eri-1 (mg366)* mutants in response to 1:100 isoamyl alcohol. $N$ = 7–10. (D) Schematic of the locomotion analysis (see Star Methods). (E–H) Normalized speed (E), overall bending activity (RMS curvature) (F), peak curvature (G), and frequency of movement (H) of wild-type animals and *eri-1 (mg366)* mutants at day 1 adult stage. $n$ = 15. Error bars indicate SEM. Unpaired t-test (B–H).

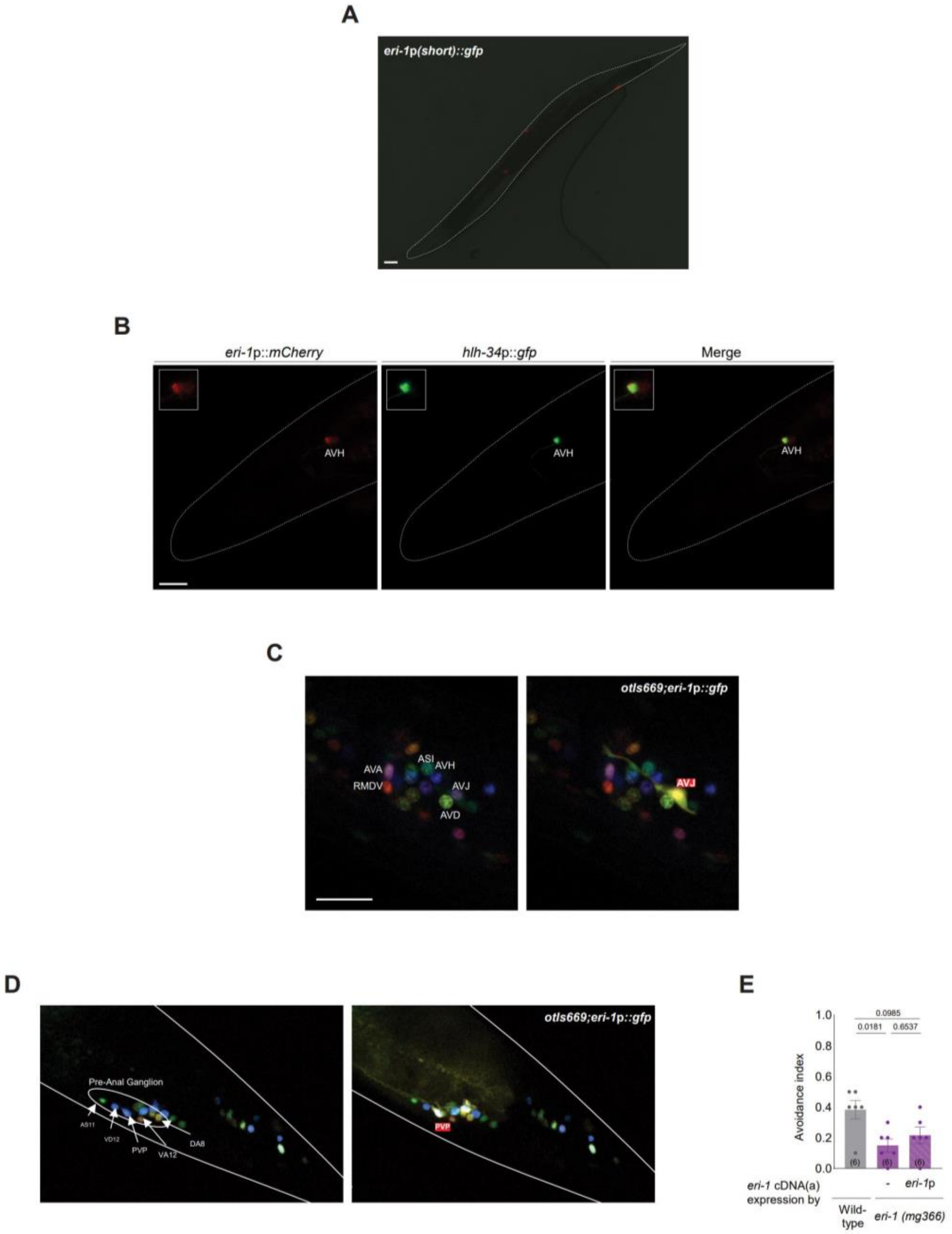


**Supplemental Figure S3.** ***eri-1* is expressed in the AVJ, AVH, and PVP neurons.** (A) Representative images of wild-type animals expressing Ex[*eri-1*p (short)::*gfp*]. The anterior is to the left. Scale bar, 20 µm. (B) Representative images of wild-type animals expressing Ex[*eri-1*p::*mCherry*] and Ex[*hlh-34p*::*gfp*]. The anterior is to the left. Scale bar, 20 µm. (C and D) Representative images of wild-type animals expressing NeuroPAL (*otIs669*) and Ex[*eri-1*p::*gfp*] and wild-type. The anterior is to the left. Scale bar, 20 µm. (E) Avoidance index of wild-type animals, *eri-1 (mg366)* mutants, and *eri-1 (mg366)* mutants expressing Ex[*eri-1*p::*eri-1* cDNA (a)] with 100 nM ascr#3. *N* = 6. Error bars indicate SEM. One-way ANOVA followed by Tukey's multiple comparisons (E).

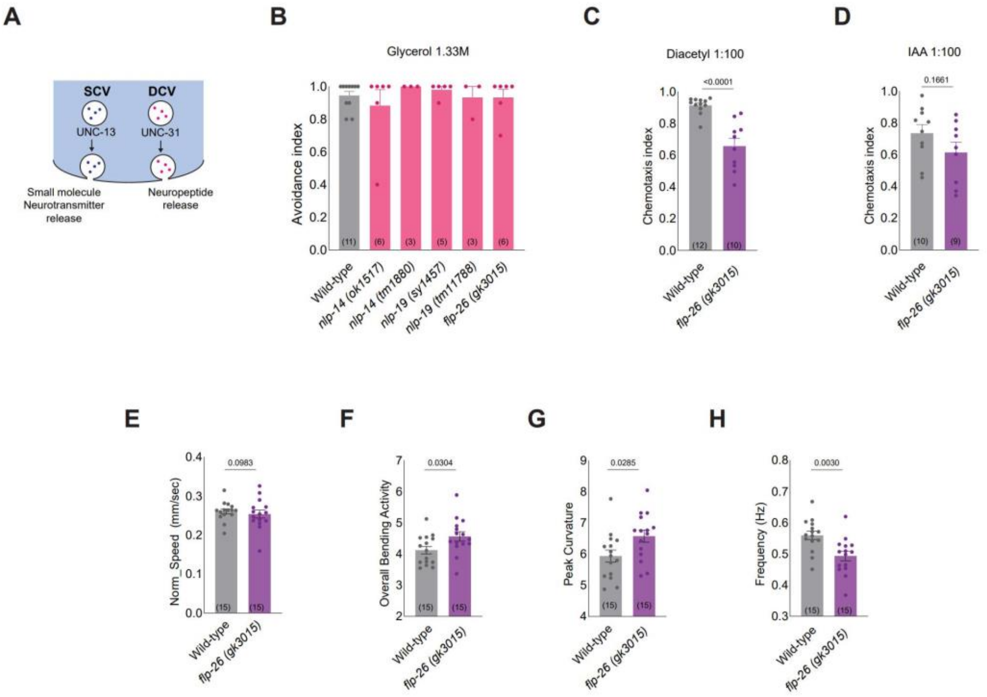


**Supplemental Figure S4.** ***flp-26* mutants retain responses to other attractants and repellents.** (A) Schematic of the roles of UNC-13 and UNC-31 in neurons. UNC-13 is required for release of small clear vesicles (SCVs) containing small molecule neurotransmitters, whereas UNC-31 mediates dense-core vesicle (DCV) exocytosis for neuropeptides. (B) Avoidance index of wild-type animals, *flp-26, nlp-14* and *nlp-19* mutants in response to 1.33 M glycerol. *N* = 3–11. (C) Chemotaxis index of wild-type animals and *flp-26 (gk3015)* mutants in response to 1:100 diacetyl. *N* = 10-12. (D) Chemotaxis index of wild-type animals and *flp-26 (gk3015)* mutants in response to 1:100 isoamyl alcohol. *N* = 9–10. (E–H) Normalized speed (E), overall bending activity (RMS curvature) (F), peak curvature (G), and frequency of movement (H) of wild-type animals and *flp-26 (gk3015)* mutants at day 1 adult stage. *n* = 15. Error bars indicate SEM. Unpaired t-test (C–H); one-way ANOVA followed by Tukey's multiple comparisons (B).

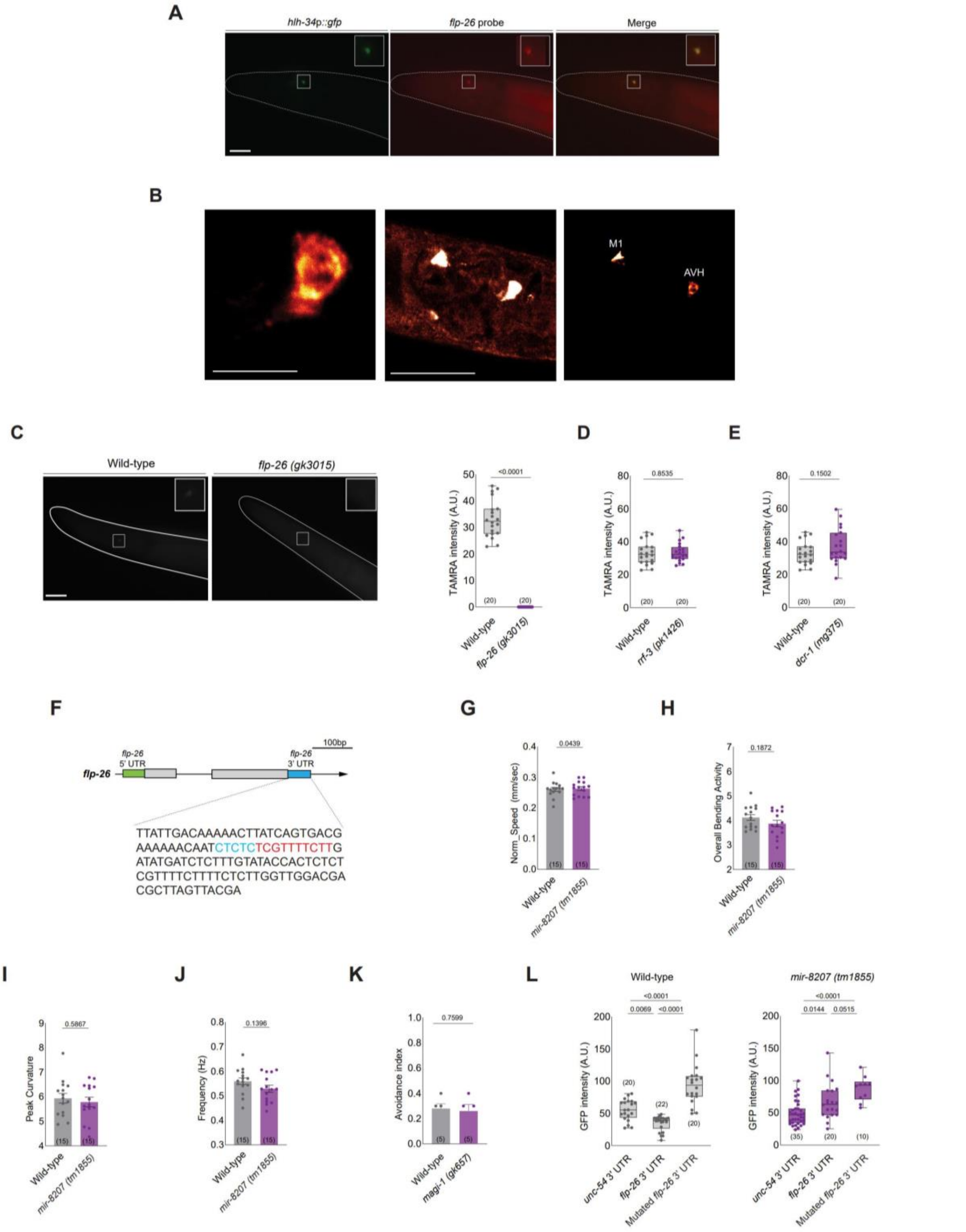


**Supplemental Figure S5. smFISH validation and genetic controls for the *eri-1–mir-8207–flp-26* axis.** (A) Representative images of smFISH transgenic animals expressing Ex[*hlh-34*p::*gfp*] labeled with the *flp-26* mRNA probe. Scale bar, 20 µm. (B) STED images of smFISH with the *flp-26* mRNA probe in wild-type animals. The AVH and M1 neurons are indicated with white arrows. Scale bar, 5 µm and 25 µm. (C) Representative images (left) and average intensity (right) of smFISH with the *flp-26* mRNA probe in wild-type animals and *flp-26 (gk 3015)* mutants. *n* = 20. Scale bar, 20 µm. (D) Average intensity of *flp-26* smFISH signal in wild-type animals and *rrf-3 (pk1426)* mutants. *n* = 20. (E) Average intensity of *flp-26* smFISH signal in animals and *dcr-1 (mg375)* mutants. *n* = 20. (F) Genomic structure and 3' UTR sequence of *flp-26* gene. Blue and red indicate predicted binding site and point mutation site, respectively. (G–J), Normalized speed (G), overall bending activity (RMS curvature) (H), peak curvature (I), and frequency of movement (J) of wild-type animals and *mir-8207 (tm1855)* mutants at day 1 adult stage. *n* = 15. (K) Avoidance index of wild-type animals and *magi-1 (gk657)* mutants in response to 100 nM ascr# 3. *N* = 5. (L) Average GFP intensity of wild-type animals and *mir-8207 (tm1855)* mutants expressing Ex[*flp-26*p::*gfp*::*unc-54* 3′ UTR], Ex[*flp-26p*::*gfp*::*flp-26* 3′ UTR], or Ex[*flp-26*p::*gfp*::*flp-26* 3′ UTR point mutation]. *n* = 10–35. Error bars indicate SEM. Unpaired t-test (C–K); one-way ANOVA followed by Tukey's multiple comparisons (L).

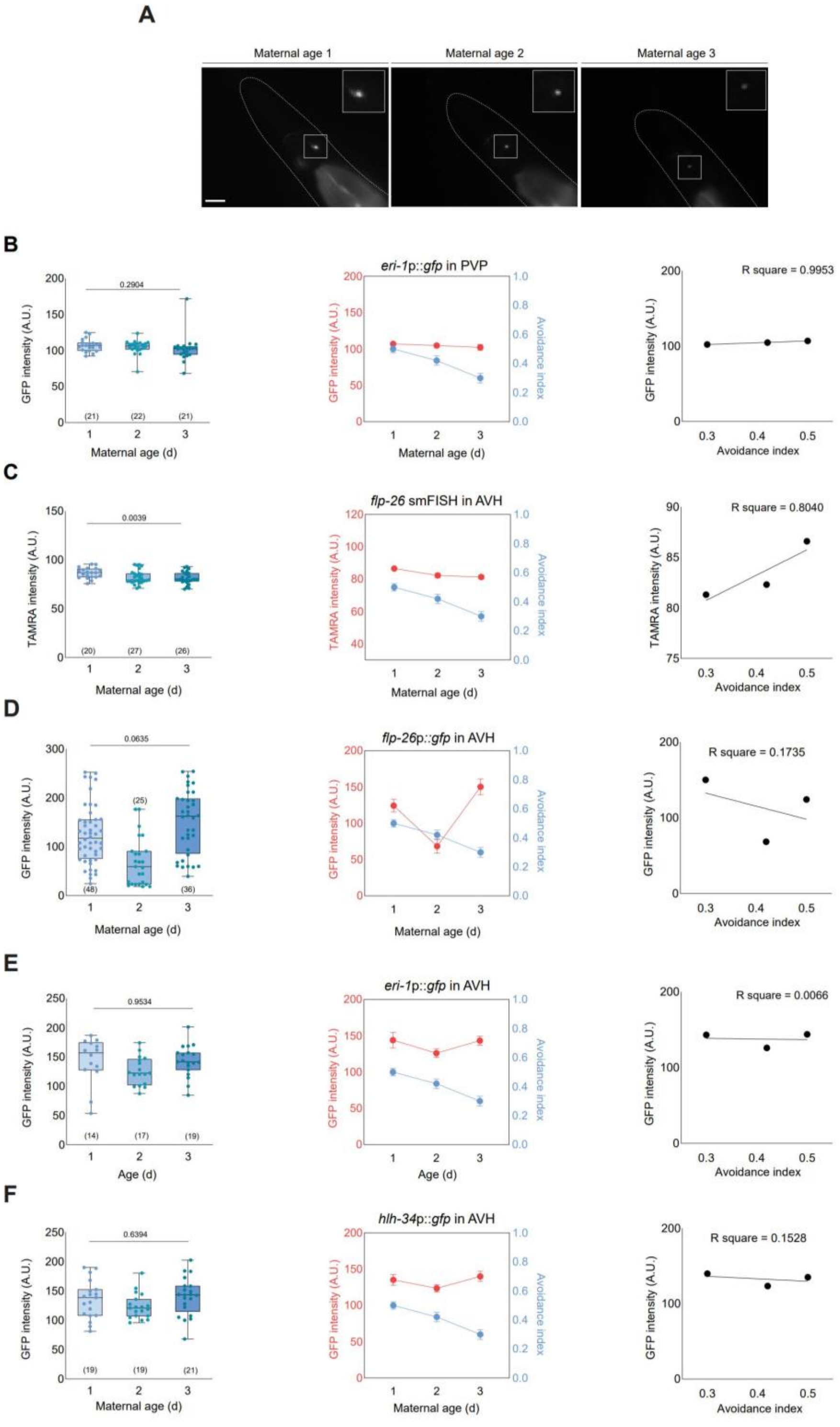


**Supplemental Figure S6. Maternal age-dependent expression of the ERI-1–*mir-8207–flp-26* pathway.** (A) Representative images of average progeny transgenic animals expressing Ex[*eri-1*p::*gfp*] from different maternal age. (B) Average GFP intensity (left), dual-axis plot with maternal age on the X-axis, GFP intensity on the left Y-axis, and avoidance index with 100 nM ascr#3 on the right Y-axis (middle), and correlation (right) of Ex[*eri-1*p::*gfp*] in PVP across maternal ages in wild-type animals. *n* = 19–29. (C) Average TAMRA intensity (left), dual-axis plot with maternal age on the X-axis, *flp-26* mRNA intensity on the left Y-axis, and avoidance index with 100 nM ascr#3 on the right Y-axis (middle), and correlation (right) of *flp-26* mRNA smFISH signal in AVH across maternal ages in wild-type animals. *n* = 20–27. (D) Average GFP intensity (left), dual-axis plot with maternal age on the X-axis, GFP intensity on the left Y-axis, and avoidance index with 100 nM ascr#3 on the right Y-axis (middle), and correlation (right) of Ex[*flp-26*p::*gfp*] in AVH across maternal ages in wild-type animals. *n* = 25–48. (E) Average GFP intensity (left), dual-axis plot with age on the X-axis, GFP intensity on the left Y-axis, and avoidance index with 100 nM ascr# 3 on the right Y-axis (middle), and correlation (right) of Ex[*eri-1*p::*gfp*] in AVH across ages in wild-type animals. *n* = 14–19. (F) Average GFP intensity (left), dual-axis plot with maternal age on the X-axis, GFP intensity on the left Y-axis, and avoidance index with 100 nM ascr#3 on the right Y-axis (middle), and correlation (right) of Ex[*hlh-34*p::*gfp*] in AVH across maternal ages in wild-type animals. *n* = 19–21. Error bars indicate SEM. Linear regression of correlation of data (B–F); one-way ANOVA followed by Tukey’s multiple comparisons (B–F).

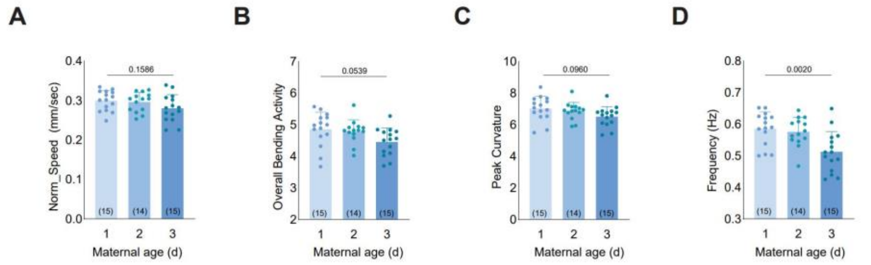


**Supplemental Figure S7. Maternal age minimally affects offspring locomotive activity.** (A–D), Normalized speed (A), overall bending activity (RMS curvature) (B), peak curvature (C), and frequency of movement (D) of wild-type progeny from different maternal age. *n* = 14–15. Error bars indicate SEM. One-way ANOVA followed by Tukey’s multiple comparisons (A–D).

**Supplementary Table 1. Neuropeptide expression level in AVH**

| | Gene name | Expression level in AVH | CENGEN expression (higher than AVH) |
|---|---|---|---|
| **FLP family** | *flp-26* | 14199.444 | M1 |
| | *flp-5* | 8003.647 | RMG, AQR, I6, FLP, AIM, PVQ, RIV, UTRY, PVT, LUA…etc |
| | *flp-12* | 2662.644 | SMB, SDQ, VA12, PDE, DA9, PDA, LUA, BDU, SAA |
| | *flp-9* | 1351.64 | RIB, RMD_LR, AVB, RIV, SMD, ADE, VB01, SIB, IL2_LR, SIA…etc |
| | *flp-28* | 792.394 | AVL, DVB, VC, HSN, M1, VC_4_5, URB, M5, PQR, RIM…etc |
| | *flp-2* | 663.823 | AIA, RID, RIB, SDQ, PQR, HSN, PVC, SAA, VC, URX…etc |
| | *flp-18* | 262.877 | RIG, AVA, VB02, DB, DB01, VA12, VB, AVE, RIV, PVN…etc |
| | *flp-32* | 226.827 | RIC, RMH, I5, URA, I6, IL2_DV, AUA, M4, PHC, URY…etc |
| **NLP family** | *nlp-50* | 40135.85 | RIP, PVM, AIY, AIM, PVD |
| | *nlp-19* | 8836.023 | N/A |
| | *nlp-14* | 6336.412 | AVF |
| | *nlp-8* | 3734.486 | ALA, AVJ, RIG, DVA |
| | *nlp-41* | 1655.275 | IL1, DVC, SMD, AIY, RIR |
| | *nlp-10* | 1577.412 | PVW, SDQ, PVT, PVR, RIF, RMG, URB, AVK, PDA, AVD…etc |
| | *nlp-21* | 997.363 | AVE, RIG, VA12, AIB, RID, RIF, AVB, DA9, AVA, AUA…etc |
| | *nlp-68* | 496.014 | M1, ADE, I5, RIP, I6, RIH, AVM, M4, I4, RMF…etc |
| | *nlp-11* | 226.854 | AIZ, PDB, URA, AIM, RIG, RIH, URB, SMD, NSM, URX…etc |

**Supplementary Table 2. List of strains used in this work**

| Genotype | Source | Strain name |
|---|---|---|
| *C. elegans* N2 | Caenorhabditis Genetics Center (CGC) | |
| *vit-2 (ac3)* | CGC | AY42 |
| *lskIs18*[*sre-1*p::GCaMP5] | CGC | KHK2532 |
| *vit-2 (ac2)*; *lskIs18*[*sre-1*p::GCaMP5] | This paper | KHK2597 |
| *vit-1 (ok2616)* | CGC | RB2565 |
| *vit-2 (ok3211)* | CGC | RB2365 |
| *vit-3 (ok2348)* | CGC | RB1815 |
| *vit-4 (ok2982)* | CGC | RB2202 |
| *vit-5 (ok3239)* | CGC | RB2382 |
| *vit-6 (tm2514)* | NBRP | - |
| *rrf-3 (pk1426)* | CGC | NL2099 |
| *dcr-1 (mg375)* | CGC | YY470 |
| *eri-1 (mg366)* | CGC | GR1373 |
| *drh-3 (ne3197)* | CGC | WM230 |
| *rde-4 (ne301)* | CGC | NE301 |
| *eri-3 (tm1361)* | NBRP | WM172 |
| *ergo-1 (tm1860)* | NBRP | WM158 |
| *alg-4 (ok1041); alg-3 (tm1155)* | CGC | WM300 |
| *alg-2 (ap426)* | CGC | PQ567 |
| *nrde-1 (gg88)* | CGC | YY160 |
| *hrde-1 (tm1200)* | NBRP | YY538 |
| *eri-1 (ok2683)* | CGC | RB2025 |
| *eri-1 (tm5288)* | NBRP | - |
| *eri-1 (tm5119)* | NBRP | - |
| *eri-1 (mg366); lskIs18*[*sre-1*p::GCaMP5] | This paper | KHK2684 |
| *lskEx1481*[*eri-1*p::*gfp, unc-122*p::*dsRed*] | This paper | KHK2083 |
| *eri-1 (mg366); lskEx1540*[*eri-1*p::*eri-1*cDNA(b), *unc-122*p::*dsRed*] | This paper | KHK2153 |
| *eri-1 (mg366)*; *lskEx1896*[*nlp-8*p::*eri-1*cDNA(b), *unc-122*p::*dsRed*] | This paper | KHK2680 |
| *eri-1 (mg366)*; *lskEx1573*[*rgef-1*p::*eri-1*cDNA(b), *unc-122*p::*dsRed*] | This paper | KHK2191 |
| *eri-1 (mg366)*; *lskEx1658*[*hlh-34*p::*eri-1*cDNA(b), *unc-122*p::*dsRed*] | This paper | KHK2331 |

| | | |
|---|---|---|
| *lskEx1658*[*hlh-34*p::*eri-1*cDNA(b), *unc-122*p::*dsRed*] | This paper | KHK2332 |
| *lskEx1788*[*hlh-34*p::HisCl1::SL2::*mCherry*, *unc-122*p::*dsRed*] | This paper | KHK2521 |
| *hlh-34 (gk1031)* | CGC | VC1904 |
| NeuroPAL *(otIs669)*; *lskEx1481*[*eri-1*p::*gfp*, *unc-122*p::*dsRed*] | This paper | KHK2582 |
| *lskEx1657*[*eri-1*p::*mCherry*, *unc-122*p::*gfp*] | This paper | KHK2328 |
| *lskEx1639*[*hlh-34p*::*gfp*, *unc-122*p::*dsRed*] | This paper | KHK2229 |
| *lskEx1657*[*eri-1*p::*mCherry*, *unc-122*p::*gfp*]; *lskEx1639*[*hlh-34*p::*gfp*, *unc-122*p::*dsRed*] | This paper | KHK2262 |
| *lskEx1572*[*eri-1*p(short)::*gfp*, *unc-122*p::*dsRed*] | This paper | KHK2189 |
| *eri-1 (mg366)*; *lskEx1540[eri-1*p::*eri-1*cDNA(a), *unc-122*p::*dsRed*] | This paper | KHK2685 |
| *unc-13 (e1091)*; *lskIs18*[*sre-1*p::GCaMP5] | This paper | KHK2688 |
| *unc-31* (*e928*); *lskIs18*[*sre-1*p::GCaMP5] | This paper | KHK2351 |
| *nlp-14 (ok1517)* | CGC | VC1108 |
| *nlp-14* (*tm1880*) | NBRP | - |
| *nlp-19 (sy1457)* | CGC | PS8682 |
| *nlp-19* (*tm11788*) | NBRP | - |
| *flp-26 (gk3015)* | CGC | VC3017 |
| *lskEx1672*[*flp-26p::mCherry, unc-122*p::*gfp*] | This paper | KHK2350 |
| *lskEx1672*[*flp-26*p::*mCherry*, *unc-122p*::*gfp*] ; *lskEx1639*[*hlh-34*p::*gfp*, *unc-122p*::*dsRed*] | This paper | KHK2351 |
| *flp-26* (*gk3015*); *lskEx1744*[*hlh-34*p::*flp-26*cDNA, *unc-122*p::*dsRed*] | This paper | KHK2540 |
| *lskEx1744*[*hlh-34*p::*flp-26*cDNA, *unc-122*p::*dsRed*] | This paper | KHK2454 |
| *flp-26* (*gk3015*); *lskIs18*[*sre-1*p::GCaMP5] | This paper | KHK2348 |
| *eri-1 (mg366*); *lskEx1671*[*flp-26*p::*gfp*, *unc-122*p::*dsRed*] | This paper | KHK2347 |
| *mir-8207 (tm1975)* | NBRP | - |
| *mir-8207 (tm1855)* | NBRP | - |
| *eri-1 (mg366); mir-8207 (tm1855)* | This paper | KHK2514 |
| *mir-8207 (tm1855)*; *lskEx1846*[*hlh-34*p::*mir-8207*, *unc-122*p::*dsRed*] | This paper | KHK2642 |
| *lskEx1671*[*flp-26*p::*gfp*, *unc-122*p::*dsRed*] | This paper | KHK2349 |
| *mir-8207* (*tm1855*); *lskEx1671*[*flp-26*p::*gfp*, *unc-122*p::*dsRed*] | This paper | KHK2517 |

| | | |
|---|---|---|
| *lskEx1798*[*flp-26*p::*gfp*::*flp-26* 3' UTR, *unc-122*p::*dsRed*] | This paper | KHK2533 |
| *mir-8207 (tm1855); lskEx1798*[*flp-26*p::*gfp*::*flp-26* 3' UTR, *unc-122*p::*dsRed*] | This paper | KHK2535 |
| *lskEx1898*[*flp-26*p::gfp::*flp-26* 3' UTR point mutation, *unc-122*p::*dsRed*] | This paper | KHK2686 |
| *mir-8207* (*tm1855*); *lskEx1898*[*flp-26*p::gfp::*flp-26* 3' UTR point mutation, *unc-122*p::*dsRed*] | This paper | KHK2687 |
| *magi-1 (gk657)* | CGC | VC1408 |
| *vit-2 (ok3211); lskEx1481*[*eri-1*p::*gfp, unc-122*p::*dsRed*] | This paper | KHK2554 |
| *lskEx1845*[*hsp*::*eri-1*cDNA(b), *unc-122*p::*dsRed*] | This paper | KHK2641 |

**Supplementary Table 3. List of oligonucleotides used in this study**

| Oligonucleotide name | Source | Cat. no. |
|---|---|---|
| *vit-2* (*ok3211*) Forward primer : ATGGAGCACGCTCTTGCTAT | CGC | N/A |
| *vit-2* (*ok3211*) Reverse primer : TGGGATCTTTCCAGAGATGG | CGC | N/A |
| *eri-1* promoter Forward primer : GCAGTTGATTGGTGTCTACGTGG | This paper | N/A |
| *eri-1* promoter Reverse primer : TCTTCTGGAGACGGCTCATCTG | This paper | N/A |
| *eri-1* cDNA(a) Forward primer : ATGAGCGCAGATGAGCCGT | This paper | N/A |
| *eri-1* cDNA(a) Reverse primer : CTAGCAAATCAAACACTTTGAATCTGTA | This paper | N/A |
| *eri-1* cDNA(b) Forward primer : ATGAGCGCAGATGAGCC | This paper | N/A |
| *eri-1* cDNA(b) Reverse primer: CTAGAAACTAGATCGATTTGAAGACG | This paper | N/A |
| *nlp-8* promoter Forward primer : TTACTGTATTCAAAATTACGGTG | This paper | N/A |
| *nlp-8* promoter Reverse primer : GAAGCTTCTGACTCATGTCGC | This paper | N/A |
| *flp-26 (gk3015)* Forward primer : TTGCTCCTCCTTCCACACTT | CGC | N/A |
| *flp-26 (gk3015)* Reverse primer : GGAAAGGGGGAAGTTGACTC | CGC | N/A |
| *flp-26* cDNA Forward primer : ATGAAGGTGATGTTCATGCTTGC | This paper | N/A |
| *flp-26* cDNA Reverse primer : CTACTTTCCAAAGCGGAGTGAAAG | This paper | N/A |
| *flp-26* 3'UTR Forward primer : TTATTGACAAAAACTTATCAGTGAC | This paper | N/A |
| *flp-26* 3'UTR Reverse primer : TTAACAAAAAAAACTTTATTAGGCA | This paper | N/A |
| *flp-26* 3' UTR site-directed mutagenesis Forward primer : TCTCTTGGTTGGACGACGCT | This paper | N/A |
| *flp-26* 3' UTR site-directed mutagenesis Reverse primer : AGTGGTATACAAGAGATCAT | This paper | N/A |
| *mir-8207 (tm1855)* Forward primer : GATTCACCTTCACATATTCCGAGC | This paper | N/A |
| *mir-8207 (tm1855)* Reverse primer : CCCAATTCGGTGGTAACAGTC | This paper | N/A |

| | | |
|---|---|---|
| *hlh-34 (gk1031)* Forward primer : GCTTTAACTCGGCCCTTAAC | This paper | N/A |
| *hlh-34 (gk1031)* Reverse primer : CGTCTTTGCTTTCTTTTCCG | This paper | N/A |
| *flp-26* promoter Forward primer : TGTGGGAAATGCGCAATGTG | This paper | N/A |
| *flp-26* promoter Reverse primer : GGAATGGAAGTCGGAAGGCA | This paper | N/A |
| *hlh-34* promoter Forward primer : GTGATACAGTTGGCTTGAGTGC | This paper | N/A |
| *hlh-34* promoter Reverse primer :GGGTTATTAAGCAACTGATG | This paper | N/A |
| *flp-26* smFISH probe 1 : GAGCAAGCATGAACATCACC | This paper | SMF-1083-5 |
| *flp-26* smFISH probe 2 : GTTGCAACGAGAGAGCTGAA | This paper | |
| *flp-26* smFISH probe 3 : TGGAATGGAAGTCGGAAGGC | This paper | |
| *flp-26* smFISH probe 4 : AGTCTTCATTAGCTCCGAAA | This paper | |
| *flp-26* smFISH probe 5 : CGTTTTGTCAATCCACTGTT | This paper | |
| *flp-26* smFISH probe 6 : TAGGGCTTCGACTCGTAGTA | This paper | |
| *flp-26* smFISH probe 7 : GTCGGCGTTAAACTCTCTTT | This paper | |
| *flp-26* smFISH probe 8 : GCTTTCCGAAACGAAGGGTT | This paper | |
| *flp-26* smFISH probe 9 : GCGAAAAAGCCAACGGTTCA | This paper | |
| *flp-26* smFISH probe 10 : TCCAAAGCGGAGTGAAAGCA | This paper | |

**Supplementary Table 4. Chemicals, commercial assays and software in this study**

| **Chemicals** | | |
|---|---|---|
| **Name** | **Source** | **Cat. no.** |
| Diacetyl | Sigma-Aldrich | Cat#8035280100 |
| Isoamyl alcohol | Sigma-Aldrich | Cat#B85307 |
| ascr#3 | Butcher lab | N/A |
| Glycerol | Sigma-Aldrich | Cat#G9012 |
| Histamine | Sigma-Aldrich | Cat#H7250 |
| Ethanol | Sigma-Aldrich | Cat#1718733 |
| Formaldehyde | Sigma-Aldrich | Cat#252549 |
| Deionized formamide | Sigma-Aldrich | Cat#S4117 |
| PBS (Phosphate-buffered saline) | Sigma-Aldrich | Cat#6506-OP |
| Ambion Nuclease-Free water | Invitrogen | Cat#AM9932 |
| **Commercial Assays** | | |
| iProof High-Fidelity DNA polymerase | BIO-RAD | Cat#1725301 |
| QIAquick PCR & Gel Cleanup Kit | QIAGEN | Cat#28506 |
| KOD FX Neo-High fidelity DNA polymerase | TOYOBO | Cat#045700 |
| Plasmid DNA Miniprep S&V Kit | BIONICS | Cat#DN10200 |
| Stellaris® RNA FISH Hybridization Buffer | Biosearch LGC | Cat#SMF-HB1-10 |
| Stellaris® RNA FISH Wash Buffer A | Biosearch LGC | Cat#SMF-WA1-60 |
| Stellaris® RNA FISH Wash Buffer B | Biosearch LGC | Cat#SMF-WB1-20 |
| **Software** | | |
| MATLAB R2023a | Mathworks | www.mathworks.com/products/matlab.html |
| GraphPad Prism 7 | Graphpad Software Inc | https://www.graphpad.com/ |
| Adobe | Adobe | https://www.adobe.com/products/illustrator.html |
| ImageJ | NIH | N/A |
| AxioVision software | Zeiss | https://www.zeiss.com/microscopy |
| Zeiss Zen (Blue Edition) Imaging Software | Zeiss | https://www.zeiss.com/microscopy |
